# Injury-related cell death and proteoglycan loss in articular cartilage: Numerical model combining necrosis, reactive oxygen species, and inflammatory cytokines

**DOI:** 10.1101/2022.06.29.498207

**Authors:** Joonas P. Kosonen, Atte S.A. Eskelinen, Gustavo A. Orozco, Petteri Nieminen, Donald D. Anderson, Alan J. Grodzinsky, Rami K. Korhonen, Petri Tanska

## Abstract

Osteoarthritis (OA) is a common musculoskeletal disease that leads to deterioration of articular cartilage, joint pain, and decreased quality of life. When OA develops after a joint injury, it is designated as post-traumatic OA (PTOA). The etiology of PTOA remains poorly understood, but it is known that proteoglycan (PG) loss, cell dysfunction, and cell death in cartilage are among the first signs of the disease. These processes, influenced by biomechanical and inflammatory stimuli, disturb the normal cell-regulated balance between tissue synthesis and degeneration. Previous computational mechanobiological models have not explicitly incorporated the cell-mediated degradation mechanisms triggered by an injury that eventually can lead to tissue-level compositional changes. Here, we developed a 2-D mechanobiological finite element model to predict necrosis, apoptosis following excessive production of reactive oxygen species (ROS), and inflammatory cytokine (interleukin-1)-driven apoptosis in cartilage explant. The resulting PG loss over 30 days was simulated. Biomechanically triggered PG degeneration, associated with cell necrosis, excessive ROS production, and cell apoptosis, was predicted to be localized near a lesion, while interleukin-1 diffusion-driven PG degeneration was manifested more globally. The numerical predictions were supported by several previous experimental findings. Furthermore, the ROS and inflammation mechanisms had longer-lasting effects (over 3 days) on the PG content than localized necrosis. Interestingly, the model also showed proteolytic activity and PG biosynthesis closer to the levels of healthy tissue when pro-inflammatory cytokines were rapidly inhibited or cleared from the culture medium, leading to partial recovery of PG content. The mechanobiological model presented here may serve as a numerical tool for assessing early cartilage degeneration mechanisms and the efficacy of interventions to mitigate PTOA progression.

**Author summary:** Osteoarthritis is one of the most common musculoskeletal diseases. When osteoarthritis develops after a joint injury, it is designated as post-traumatic osteoarthritis. A defining feature of osteoarthritis is degeneration of articular cartilage, which is partly driven by cartilage cells after joint injury, and further accelerated by inflammation. The degeneration triggered by these biomechanical and biochemical mechanisms is currently irreversible. Thus, early prevention/mitigation of disease progression is a key to avoiding PTOA. Prior computational models have been developed to provide insights into the complex mechanisms of cartilage degradation, but they rarely include cell-level cartilage degeneration mechanisms. Here, we present a novel approach to simulate how the early post-traumatic biomechanical and inflammatory effects on cartilage cells eventually influence tissue composition. Our model includes the key regulators of early post-traumatic osteoarthritis: chondral lesions, cell death, reactive oxygen species, and inflammatory cytokines. The model is supported by several experimental explant culture findings. Interestingly, we found that when post-injury inflammation is mitigated, cartilage composition can partially recover. We suggest that mechanobiological models including cell–tissue-level mechanisms can serve as future tools for evaluating high-risk lesions and developing new intervention strategies.

## 1. Introduction

Joint injuries trigger cell biological signaling pathways implicated in articular cartilage degeneration [1–3]. Cartilage has a limited innate capacity for repair, so when joint injuries cause loss of chondrocyte (cartilage cell) viability and extracellular matrix (ECM) components, it can be irreversible. Ultimately, these sequelae of joint injury lead to post-traumatic osteoarthritis (PTOA), a disease marked by pain in the affected joint [1, 2]. The mechanisms of the onset and progression of PTOA are poorly understood, but several intertwined factors have been identified: chondrocyte death [4, 5], mitochondrial dysfunction and the subsequent overproduction of reactive oxygen species (ROS) [6, 7], increased proteolytic activity triggered by excessive mechanical loading [8], and inflammation [2].

Mechanical loading is an important factor in chondrocyte-regulated cartilage homeostasis [9, 10]. Injurious loading may initiate ECM degeneration [1,7,11] and cause cell death including apoptosis and necrosis [10,12–15]. This degenerative pathway may be further promoted locally by dynamic loading, even if compressive tissue-level mechanical strains remain within physiological limits [16]. Necrosis is an acute form of cell death caused by direct mechanical damage to cells such as injurious single-impact loading or high local strains and/or strain rates [10,12,13,17]. Necrosis is also suggested to result in the release of damage-associated molecular patterns (DAMPs) and pro-inflammatory cytokines [18–20] and lead to ECM degeneration caused by proteolytic enzymes [21]. In addition, near the injury site, excessive local strains may alter cell function. For instance, associated changes in mitochondrial activity and physiology can culminate in the excessive production of ROS [22, 23]. Apoptosis, the controlled subacute form of cell death, has also been associated with excessive production of ROS [14, 24]. Excessive ROS production has been suggested to promote ECM degeneration via decreased matrix biosynthesis [25], increased release of proteolytic enzymes [26, 27], and inhibition of tissue inhibitors of metalloproteinases (TIMPs) [25, 28].

Inflammation is another important factor in cartilage homeostasis. During the early phases of PTOA, the introduction of pro-inflammatory cytokines such as interleukin-1 (IL-1), IL-6, IL-18, and tumor necrosis factor-α (TNF-α) predisposes cartilage to degeneration [2,29,30]. Catabolism and reduced biosynthesis in the ECM is counter-balanced with anti-inflammatory cytokines (*e.g.*, IL-4, IL-10, IL-13) [30], TIMPs [31], and growth factors such as insulin-like growth factor-1 [29, 30]. Prolonged inflammation may shift cartilage homeostasis to the catabolic state, in which the ECM is degraded [2, 32] via aggrecanases (*e.g.*, disintegrin and metalloproteinase with thrombospondin motifs-4,5; ADAMTS-4,5) and collagenases (*e.g.*, matrix metalloproteinases-1,3,13; MMP-1,3,13) [2,30,31].

The ability to predict cartilage degeneration via both biomechanical and inflammatory mechanisms is critical to comprehending disease progression, evaluating the efficacy of medical treatments, and developing new intervention strategies. Computational modeling has great potential in this regard while being cost-efficient. Previous computational finite element models have introduced promising frameworks to simulate the biomechanics- and inflammation-driven cartilage degeneration at joint, tissue, and cell levels in both a spatial and temporal manner [16,33–36]. Previous biomechanics-driven computational models have targeted primary cartilage injury mechanisms including necrosis, apoptosis, and pro-inflammatory cytokine and DAMP-signaling without including the degeneration of different ECM components [35,37,38]. More recently, strain/stress threshold-based modeling approaches have been developed to predict tissue-level proteoglycan (PG) loss without explicitly modeling the underlying chondrocyte-regulated mechanisms [33, 39]. No previous computational approach has modeled the chondrocyte-driven biomechanical and biochemical mechanisms triggered by injury and regulating spatial and temporal tissue-level degeneration.

In this study, we developed a new 2-D cell-and-tissue-level mechanobiological model of cartilage degeneration [16,34,36] to predict injury-related cell responses after excessive biomechanical loading, inflammation, and subsequent early-stage PTOA progression. We did not model the injurious loading *per se*, but we instead concentrated on how cell death and compositional changes evolve in injured cartilage that is possibly experiencing locally elevated strains post-injury. We hypothesized that i) injury-related catabolic mechanisms (necrosis and apoptosis) and PG loss occur at early time-points in close proximity to lesions while ii) inflammation-mediated PG loss occurs later and in more distant intact areas. To predict tissue-level cell death and PG loss in an injured environment, we simulated three different cell-level mechanisms separately and simultaneously. In the numerical model, excessive biomechanical shear strains trigger i) necrosis and ii) apoptosis following cell damage (e.g., mitochondrial dysfunction) and ROS overproduction, while IL-1 diffusing into the tissue trigger iii) inflammatory responses. We compared the simulated cell death and PG content predictions with outcomes of *in vitro* experiments (15,40). To address the lack of quantitative experimental data, we conducted a sensitivity analysis for the most relevant parameters in the model, which were selected based on preliminary simulations (necrosis/cell damage rate, ROS production rate, rate of spontaneous apoptosis, and decay rate of IL-1 concentration). Our approach is a novel step towards modeling PTOA progression through chondrocyte-driven biological mechanisms triggered by both locally excessive biomechanical loading and inflammation.

## 2. Materials and methods

A computational mechanobiological model, inspired by previous models [16,34,36,37], was developed to simulate cartilage degeneration in experimental cartilage geometry after injurious unconfined compression to explain biological tissue-level damage via cell-driven mechanisms [16, 40]. The cartilage PG degeneration was controlled with three different adaptive mechanisms (Fig. 1): shear strain-induced A) necrosis of a cell population and B) ROS overproduction by remaining live cells, which further leads to cell apoptosis. These injury-related mechanisms ultimately resulted in an increased aggrecanase release from regions containing ROS-affected cells. The last mechanism is associated with the effects of IL-1, which can cause chondrocyte apoptosis as well as upregulation of aggrecanase in the remaining live cells. All three mechanisms were assumed to lead to decreased PG biosynthesis and were modeled separately and also simultaneously in a combined model. We simulated the evolution of the viable cell and matrix PG content distributions for 12 days, while also providing extrapolated insights up to 30 days. Based on the simulated results, we quantitatively analyzed near-lesion (0.1 mm from lesion edge) and bulk cell viability and PG loss at several time-points. The simulated results in an injured cartilage explant model were also qualitatively compared with previous explant culture experiments.

**Fig 1.**
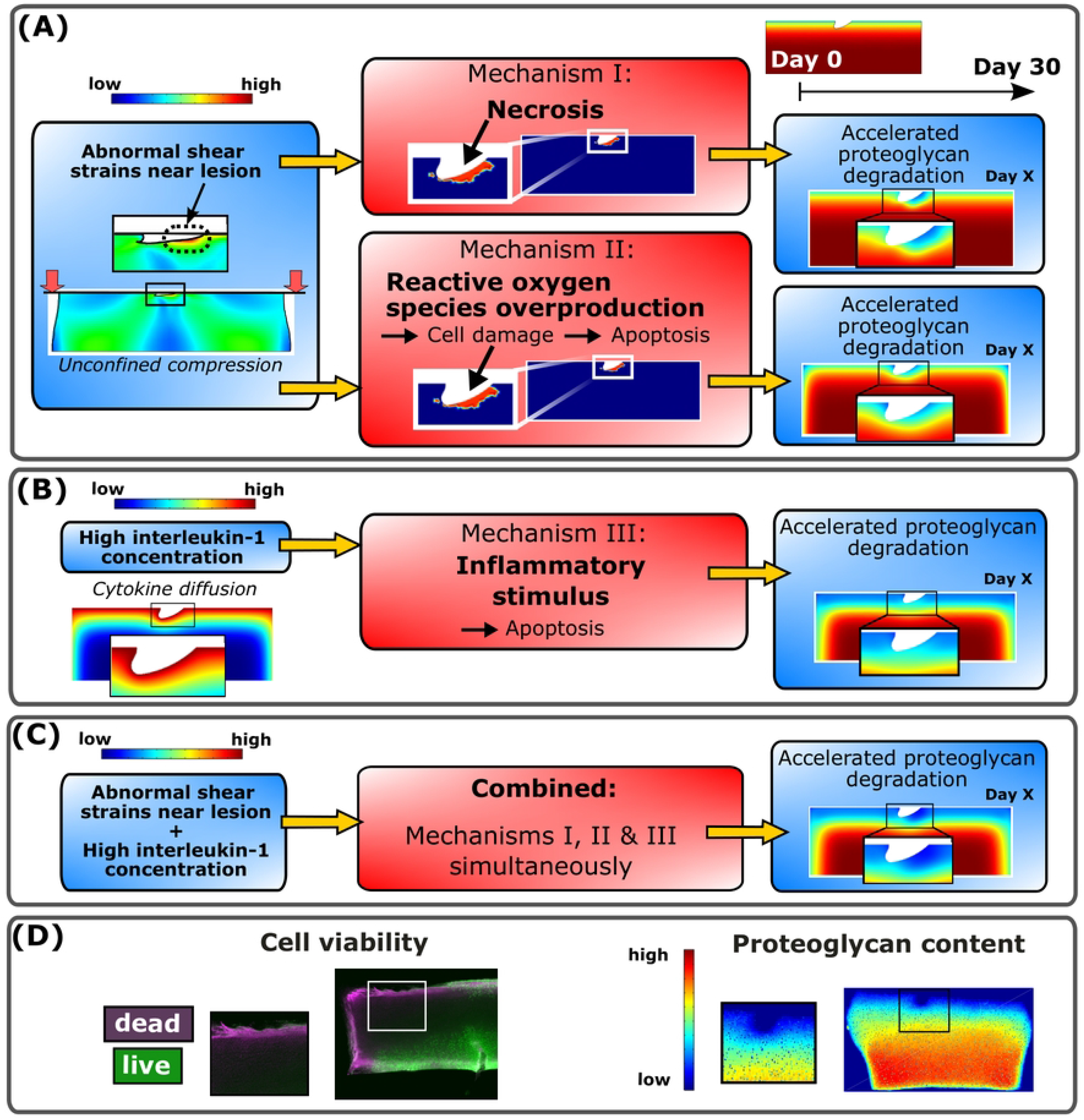
Computational modeling framework and comparison against biological data. Delineation of the simulated mechanisms I-III in the proposed computational model to predict temporal and spatial changes in cell viability and proteoglycan (PG) loss for 30 days. (A) Unconfined compression (15% axial strain, 1 Hz loading frequency) of injured cartilage was simulated to obtain maximum shear strain distributions. In areas experiencing abnormal maximum shear strains, two biomechanically-induced degradation were triggered; chondrocyte necrosis (mechanism I) and cell damage reactive oxygen species overproduction followed by apoptosis (mechanism II). (B) Interleukin-1 (IL-1) diffusion (1ng/ml of IL-1 in the culture medium) causing high spatially distributed IL-1 concentration in the cartilage caused inflammatory cell stimulus. This led to chondrocyte apoptosis (mechanism III). Moreover, all the mechanisms I-III accelerated the proteoglycan degradation by decreasing the PG biosynthesis and increasing the proteolysis of PGs. (C) Finally, combined model was developed to simulate the sygergistic effects of mechanisms I-III. (D) Simulated cell viability and proteoglycan content were also generally compared against experimentally measured cell viability and digital densitometry measurements (∼proteoglycan content).

### 2.1. Comparative biological data

Predictions of our theoretical computational model were visually compared against histological changes observed in the previous explant culture experiments (Fig. 2) [16, 40]. We highlight that the exact experimental protocol was not modeled, thus no quantitative comparison is provided. We find this visual comparison feasible, since the goal in this study was to gain understanding of the possible underlying mechanisms to explain experimental findings in PTOA-like conditions.

**Fig 2.**
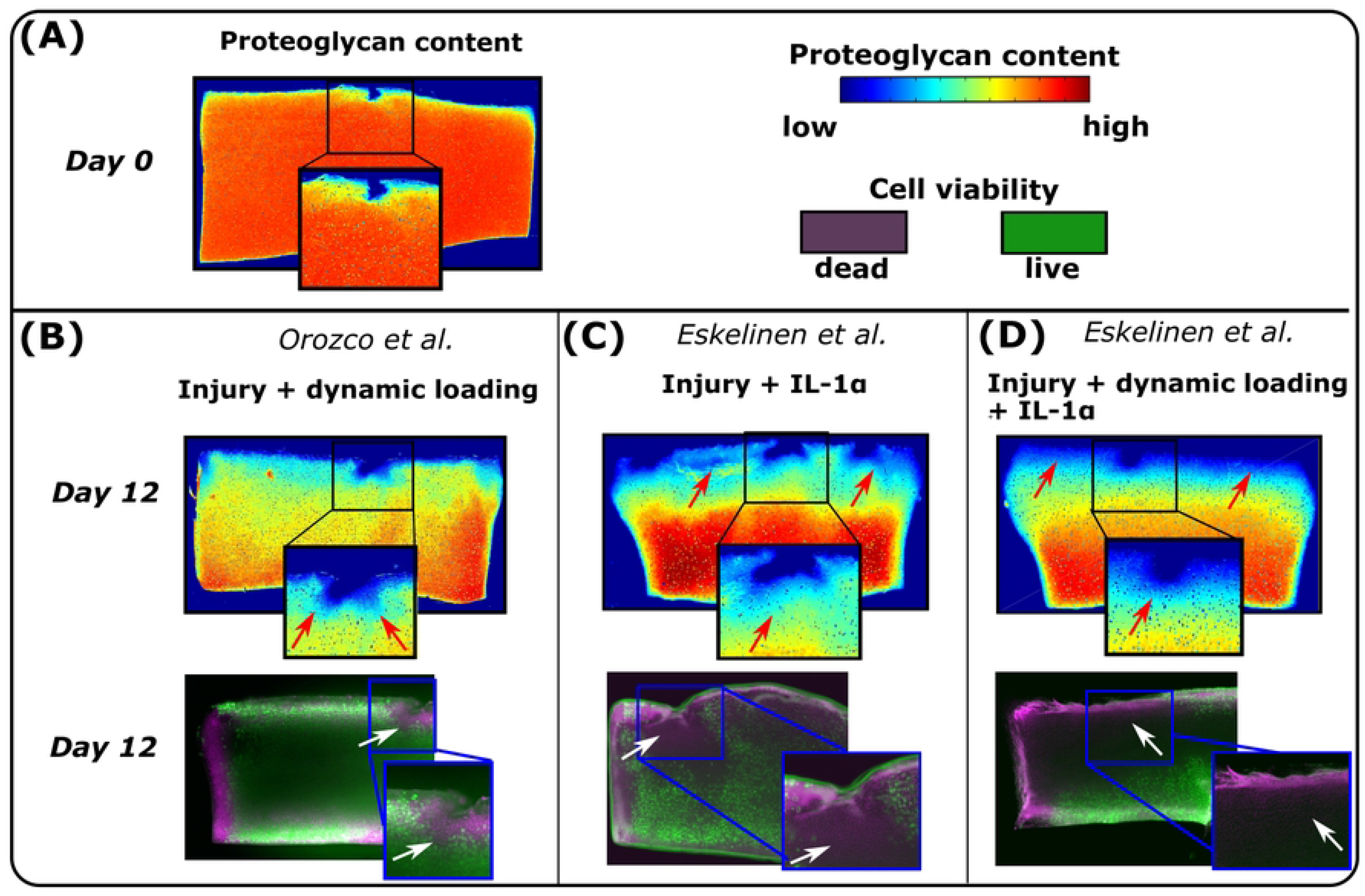
Previous experiments. In the previous experiments conducted by Orozco et al [16] and Eskelinen et al. (40), the injured, dynamically loaded and IL-1 challenged cartilage samples were analyzed at several time-points during 12-day culture. Cell viability and proteoglycan content (∼optical density) were measured with fluorescence microscopy and digital densitometry, respectively. (A) At day 0, proteoglycan loss in cartilage was minor. At day 12, the results showed (B) substantial cell death and proteoglycan loss near lesion after dynamic loading in the injured cartilage. Interleukin-1 challenge induced cell death and PG loss also in the intact areas (C) with and (D) without dynamic loading post-injury. Red arrows highlight locally decreased optical density and white arrows increased local cell death.

In the experiments (Fig. 2) [16, 40], cylindrical articular cartilage samples (diameter 3 mm, thickness 1 mm) were prepared from patellofemoral grooves of freshly slaughtered 1-2-week-old calves. The samples were subjected to injurious compression (50% strain, 100%/s strain rate) with 1) compressive cyclic loading (15% strain amplitude, 1 Hz haversine waveform, 1 hour loading periods 4 times per day) 2) IL-1-challenge (1 ng/ml), or 3) a combination of compressive cyclic loading and IL-1 challenge. Injurious compression resulted in a physical cartilage lesion in all samples. A free-swelling control group was also included for comparison. Cell viability and localized PG content were assessed at several timepoints up to 12 days with fluorescence microscopy and digital densitometry.

The experiments [16, 40] showed minor cell death and PG loss (decreased optical density) between intact and injured sample regions on the day of injury (Fig. 2A, day 0). Qualitatively, the PG content in the injured and dynamically loaded group decreased mostly near the lesion (Fig. 2B, day 12 vs. day 0, red arrows). After injury and IL-1 treatment, PG content decreased noticeably near all edges of the cartilage plug (Fig. 2C, red arrows). Dynamically loaded injured and inflamed plugs also experienced marked PG loss both away and near the lesion (Fig 2D).

### 2.2. Simulation of abnormal biomechanical shear strains promoting necrosis and cell damage

A finite element model of injured cartilage was subjected to physiologically relevant dynamic loading as in a previous study [16]. The injury (lesion) and simulated dynamic loading (two unconfined compressions) were implemented based on the experiments [16]. Importantly, we did not model the injurious loading itself, but rather the subsequent physiologically relevant dynamic loading of injured cartilage. The mechanical behavior of cartilage was modeled using a fibril-reinforced porohyperelastic material with Donnan osmotic swelling [41]. The material model input incorporated depth-dependent material properties including water content, PG content, and collagen orientation and density [16] (Supplementary material section S1). This material model has been shown to reliably capture cartilage mechanical behavior [41, 42]. The model output was maximum shear strain distribution, showing locally elevated shear strains near the lesions, even though tissue-level loading remained within physiological limits [16, 39] (Fig. 1A). The mechanical model was constructed in ABAQUS (v. 2021, Dassault Systèmes, Providence, RI, USA), and solutions were obtained using ‘soil consolidation’ analysis (transient analysis of partially or fully saturated fluid-filled porous media) with the same model geometry and finite element mesh that was assured to converge in our previous work (918 linear axisymmetric elements with pore pressure, element type: CPE4P) [16]. Boundary conditions were assigned as in the previous model (Supplementary material section S2). Since excessive shear strains cause cell death in cartilage [17], we used the maximum shear strain distribution as a driving parameter for the locally triggered cell death and PG loss. As a preliminary test, we conducted simulations with higher compressive strain amplitude to estimate areas experiencing cell necrosis/damage triggered after dynamic high-strain tissue level compression (40% unconfined axial compressions, 1 Hz loading frequency). For more detailed information readers are referred to Supplementary material section S3.

### 2.3 Modeling cell death and PG loss

#### Diffusion of aggrecanases and decrease in PG biosynthesis

Injury-related cell death and damage, as well as diffusing inflammatory cytokines, may lead to release of aggrecanases [8, 21]. In our model, mechanisms I–III (see below) regulated the amount of released aggrecanases diffusing in cartilage and suppressed PG biosynthesis after decreased cell viability, both leading to PG loss. Also, PGs may diffuse out of the tissue passively through the cartilage–fluid-interface. These mechanisms were modeled with time-dependent reaction–diffusion partial differential equations [36]

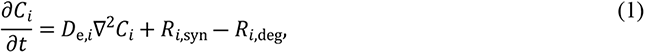

where *t* is time, *C*_*i*_ is the concentration of the biochemical species *i* (aggrecanases, PGs, IL-1, viable/necrotic/damaged cells), *D*_e,*i*_ is the effective diffusivity (zero for cell populations, as we assumed no cell migration), *R*_*i*,syn_ is the source (synthesis) term, and *R*_*i*,deg_ is the sink (degeneration) term of the species *i*. The source/sink terms utilized Michaelis–Menten kinetics like the model by Kar et al. [36]. For example, an increase in the aggrecanase concentration increases the PG sink term, whereas cell death decreases the PG source term. The initial PG content was obtained from the previous experiments [36, 43]. For more detailed information, readers are referred to Supplementary Material section S4. Diffusion and reaction of species *i* were modeled in COMSOL Multiphysics (version 5.6, Burlington, MA, USA) using a 2405-element triangular mesh (Fig. S5 Supplementary Material section S5).

#### Mechanism I. Necrosis

First, regions presumed to experience early necrosis due to high mechanical strain [13,17,44] were obtained from ABAQUS simulations using a custom-written (Supplementary Material section S6) MATLAB script (R2018b, The MathWorks, Inc., Natick, MA, USA). Based on earlier studies, we assumed that when the maximum shear strain in an element exceeded a threshold of 50% [16], 40% of cells were assumed to become necrotic [45]. These live and necrotic cell distributions were then imported into COMSOL.

The presence of necrotic cells was assumed to result in a rapid increase of local aggrecanase concentration. The imported necrotic cell distribution then served as an initial condition for the enzymatic (aggrecanase-induced) PG degradation. Acute necrosis-driven PG degeneration via aggrecanases is supported by experimental findings reporting rapid cell death within hours after single-impact loading [45] and studies suggesting necrosis-driven release or stimulation of proteolytic enzymes [21]. According to our preliminary tests, this choice also showed similarities with experimentally observed early cell death and PG loss near cartilage lesions [16, 40]. In addition, it has been suggested that high local strains during repetitive dynamic loading in injured cartilage could lead to accumulated cell death and possibly secondary necrosis in the superficial zone [46, 47], promoting the localized release of inflammatory factors [19–21] which could increase the proteolytic activity associated with the surviving cells [30]. Thus, we assumed an acute aggrecanase release (concentration *C*_aga,init_) from necrotic cells *C*_n,c_ at the beginning of the simulation:

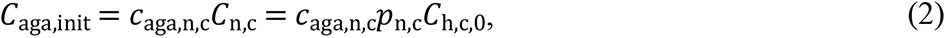

where *c*_aga,n,c_ is a calibration constant for the released aggrecanase (1.2 · 10^―19^ mol) based on a visual comparison of simulated PG concentration and histologically observed PG content findings [40], *p*_n,c_ = 0.4 = 40% is the fraction of necrotic cells [45], and 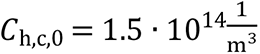 is the initial concentration of healthy cells [48].

#### Mechanism II. Damaged cells, ROS release, and apoptosis

Similarly as with necrosis, we assumed that 40% of the cells experiencing the maximum shear strains > 50% will become ‘damaged cells’ *C*_d,c_ (e.g., experiencing mitochondrial dysfunction) [16]:

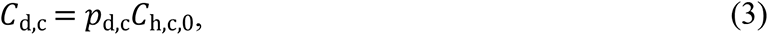

where *p*_d,c_ = 0.4 = 40% is the fraction of damaged cells [49]. Based on observations of increased ROS production in response to excessive mechanical loading [14,23,49], we assumed that the localized ROS concentration *C*_ROS_increases as a function of damaged cell concentration *C*_d,c_ [37]:

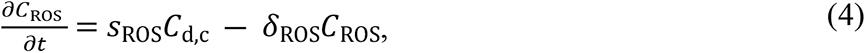

where 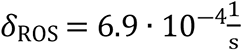 is the ROS decay rate [37] and *s*_ROS_ is the ROS synthesis rate described as

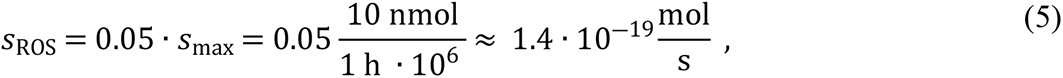

where *s*_max_ is the estimated maximum oxygen consumption rate (5–21% oxygen tension) [37, 50]. Moreover, since the ROS production in healthy cartilage has been estimated to be 1–3% of the maximum oxygen consumption [24,37,51], we assumed 5% ROS production in injured cartilage (overproduction). Moreover, we assumed no diffusion of ROS since the approximate half-life of the mitochondrial ROS is relatively short (< 1 ms) [52]. Excessive ROS production has been suggested to result in apoptosis and PG loss [14,53,54]. The former phenomenon was incorporated as damaged cells *C*_d,c_ turning apoptotic in an exponential manner [55, 56]:

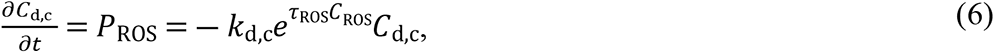

where *P*_ROS_ describes the rate of damaged cells turning apoptotic due to 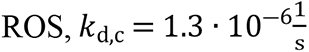 is cell death rate for damaged cells [57], and *τ*_ROS_ a calibration coefficient for ROS-dependent cell death 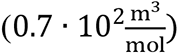. Furthermore, PG loss was affected by increased aggrecanase release due to ROS, modulated by a modified stimulus equation originally introduced by Kar et al. [36] (Supplementary Material section S3). Finally, PG degeneration was modeled based on Eq. (1).

#### Mechanism III. Inflammation-induced apoptosis

Pro-inflammatory cytokine-mediated apoptosis was implemented with IL-1 in the following exponential equation [58]

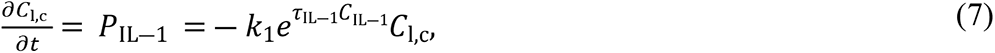

where *C*_l,c_ is the concentration of live cells (*C*_l,c_ = *C*_h,c,0_, if only inflammation is considered or *C*_l,c_ = *C*_h,c,0_ (1 ― *p*_n,c_ ― *p*_d,c_) if also necrosis and cell damage are considered in the cells experiencing over 50% maximum shear strain), 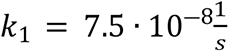 is the rate of spontaneous apoptosis (11 % of cells are apoptotic after 17 days under free-swelling conditions without exogenous cytokines) [29], 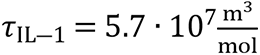 is a calibration coefficient for experimentally observed IL-1-induced depth-dependent apoptosis [29], and *C*_IL―1_is IL-1 concentration. The chosen IL-1 concentration was 1 ng/ml, implemented as a Dirichlet boundary condition on all the edges except the bottom of the cartilage geometry [29, 36]. Cytokine diffusion led to PG loss after loss of cell viability and upregulation of aggrecanases via IL-1-mediated stimulus which were simulated separately and simultaneously (See Supplementary Material section S4).

#### Combining injury-related and inflammatory mechanisms

In the combined model, cell death including injury-related i) necrosis, ii) apoptosis via ROS overproduction in the damaged cells, and iii) IL-1-induced apoptosis were all considered simultaneously. Here, the live cell concentration was affected as described in Eq. (7). The damaged cells could turn apoptotic due to ROS overproduction (*P*_ROS_, Eq (6)) and inflammation (*P*_IL―1_, Eq (7)).

### 2.4 Sensitivity analysis for the computational model parameters

To address a lack of experimental data needed to calibrate some model parameters, we conducted a computational sensitivity analysis for the essential parameters affecting cell death and PG loss. Based on our preliminary tests during model development, the chosen parameters were necrosis fraction (*p*_n,c_), damaged cell fraction (*p*_d,c_), ROS production rate (*s*_ROS_, healthy and excessive levels), and rate of spontaneous apoptosis (*k*_1_; the IL-1-induced aggrecanase stimulus was turned off to perceive the effect of altered PG biosynthesis due to cell death on PG loss; Table 1).

**Table 1.**
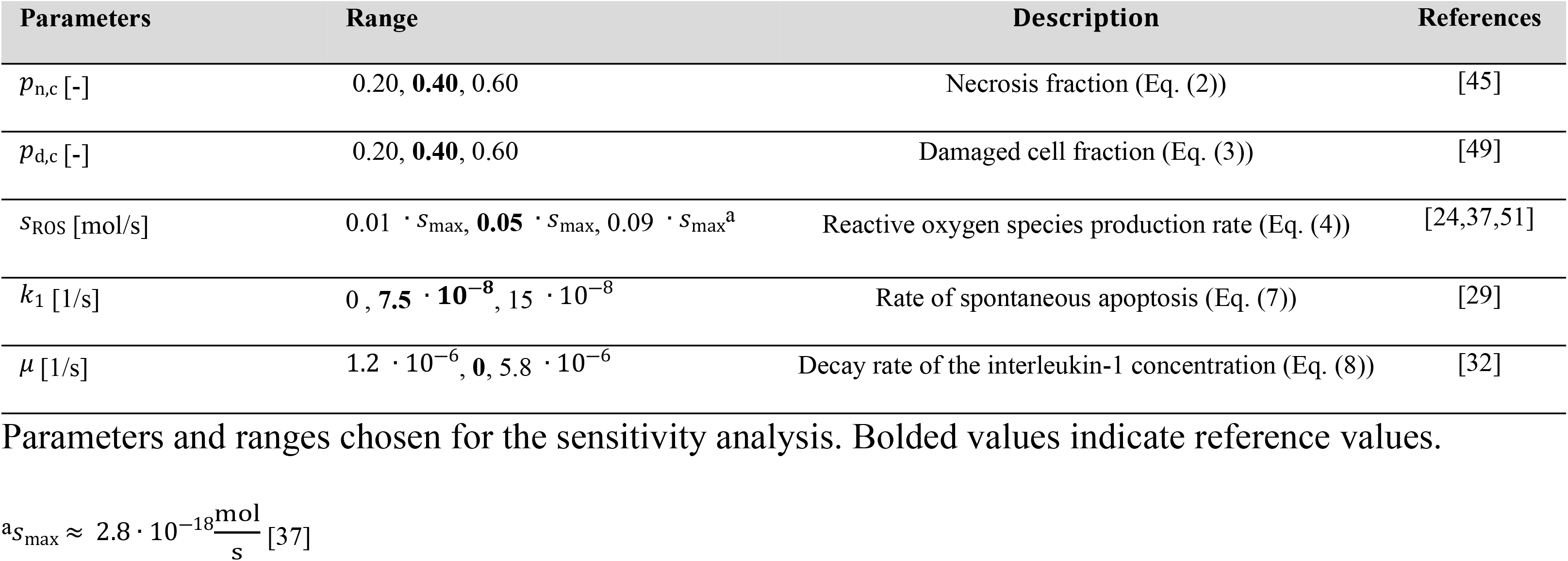
Parameters for sensitivity analysis.

#### Decreased IL-1 concentration

Previous clinical and pre-clinical studies have suggested that inflammation may play a major role in PTOA progression driven by inflammatory cytokines, but after acute inflammation, the concentration of the pro-inflammatory cytokines can decrease exponentially [32, 59]. Hence, to gain insights into the possible resolution of acute inflammation and tissue recovery, we simulated time-dependent slow and fast exponential decreases of IL-1 concentration in the culture medium as

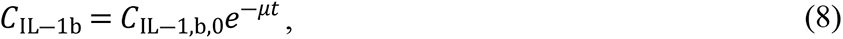

where *C*_IL―1,b,0_ is the initial boundary concentration of IL-1 (1 ng/ml) and *μ* is the decay rate of the IL-1 concentration.

## 3. Results

### 3.1. Necrosis

In the numerical simulations, necrotic cell death was localized near the cartilage injury (Fig. 1A). At day 5, the computational reference model (*p*_n,c_ = 0.4) predicted that 10.8% of the viable cells would be necrotic and 21.6% of PGs would be lost within 0.1 mm from the cartilage lesion compared to day 0 (Figs. 3A and 4, red line). The simulated PG content decreased rapidly and locally during the first day, followed by partial recovery for the rest of the simulation. Sensitivity analysis revealed that, at day 5, a smaller number of necrotic cells (*p*_n,c_ = 0.2; Fig. 4B, blue line) resulted in an average PG loss of 16.4% while a greater number (Fig. 4C, blue line) of necrotic cells (*p*_n,c_ = 0.6; Fig. 4B, purple line) resulted in an average PG loss of 26.1% (Fig. 4D, purple line).

**Fig 3.**
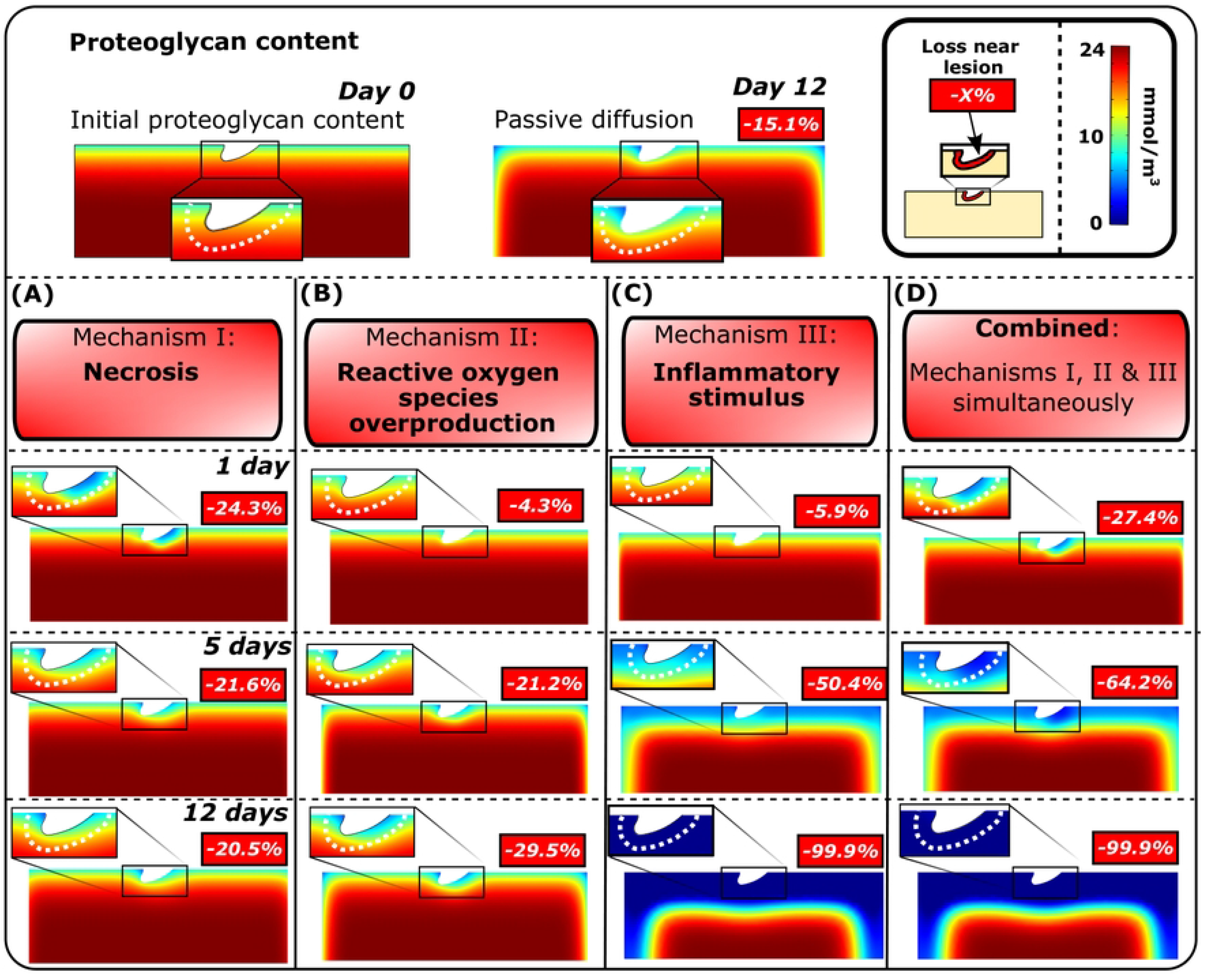
Simulated proteoglycan degeneration. Comparison of the simulated spatial changes in proteoglycan content after A) acute necrosis, B) cell damage, subsequent overproduction of reactive oxygen species and apoptosis, C) inflammatory stimulus, and D) combined mechanisms I, II and III at days 1, 5 and 12 showed different temporal changes in proteoglycan distribution. Percentual changes in the proximity of the simulated lesion (0.1 mm from lesion edge) are computed relative to proteoglycan content at day 0.

**Fig 4.**
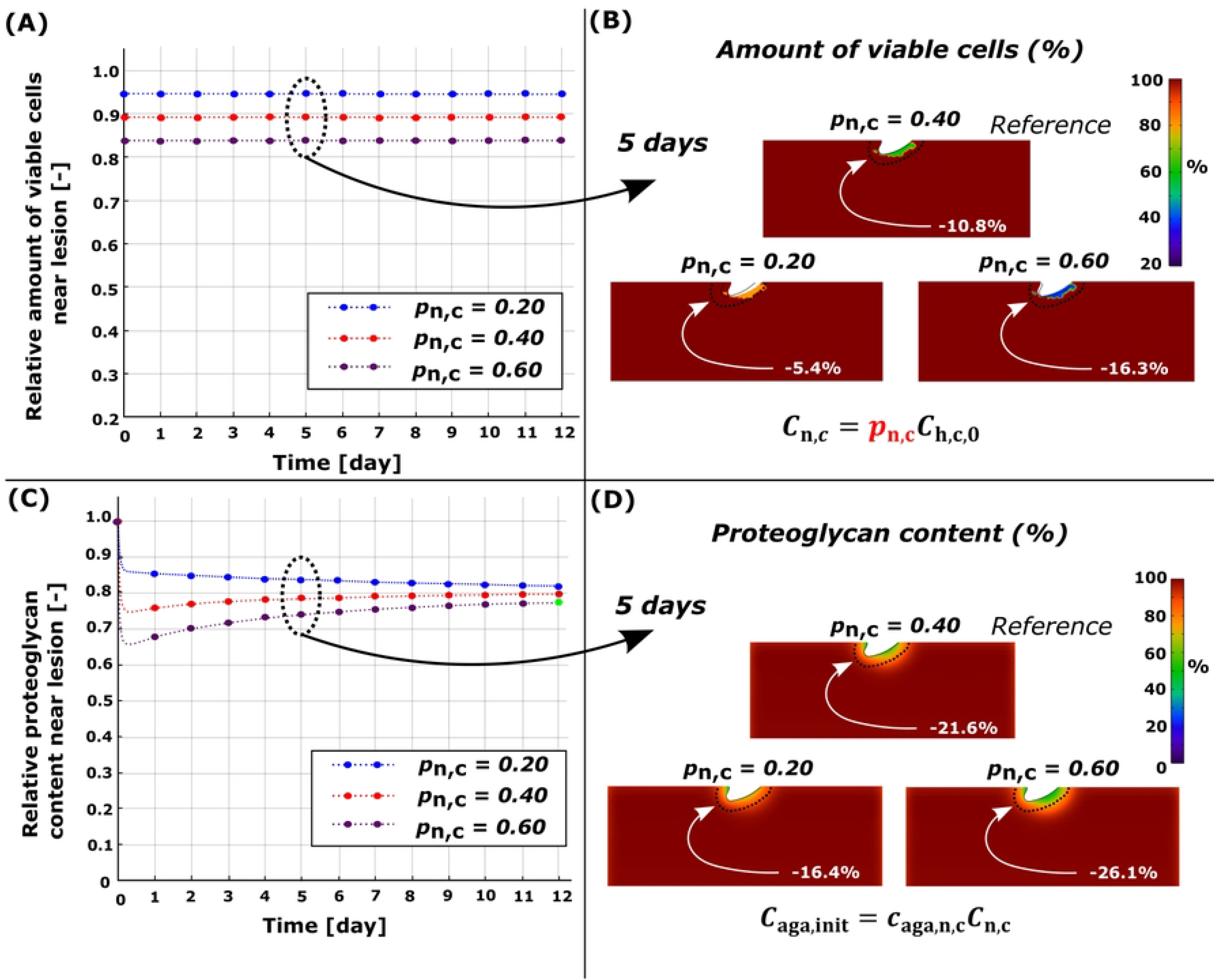
Sensitivity analysis of simulated necrosis rate *p*_n,c_. Comparison of temporal changes and spatial changes at day 5 in (A)-(B) in cell viability and (C)-(D) in proteoglycan content. (C) Higher necrosis rate led to fast proteoglycan degeneration at early time-points (days 0-1) and partial recovery proteoglycan content (days 0-3) near the cartilage lesion. Red line in (A) and (C) and refers to the reference model (*p*_n,c_ = 0.40).

### 3.2. Damaged cells, ROS release, and apoptosis

Cell damage was observed also near the lesion (Fig. 1A). The computational reference model (moderate ROS overproduction) showed average cell apoptosis of 6.5% and average PG loss of 21.2% near the lesion at day 5 (Figs. 3B and 5, red line). An 80% decrease in ROS production rate (low, healthy levels; Fig. 5, blue line) showed simulated apoptosis of 5.0% and PG loss of 13.0%, whereas increasing ROS production (high ROS overproduction; Fig. 5, purple line) to excessive levels led to apoptosis of 7.5% and PG loss of 26.4%. Changes in the damaged cell fraction showed a similar effect on the apoptosis and PG content compared to variations in the ROS production rate (Fig. 6).

**Fig 5.**
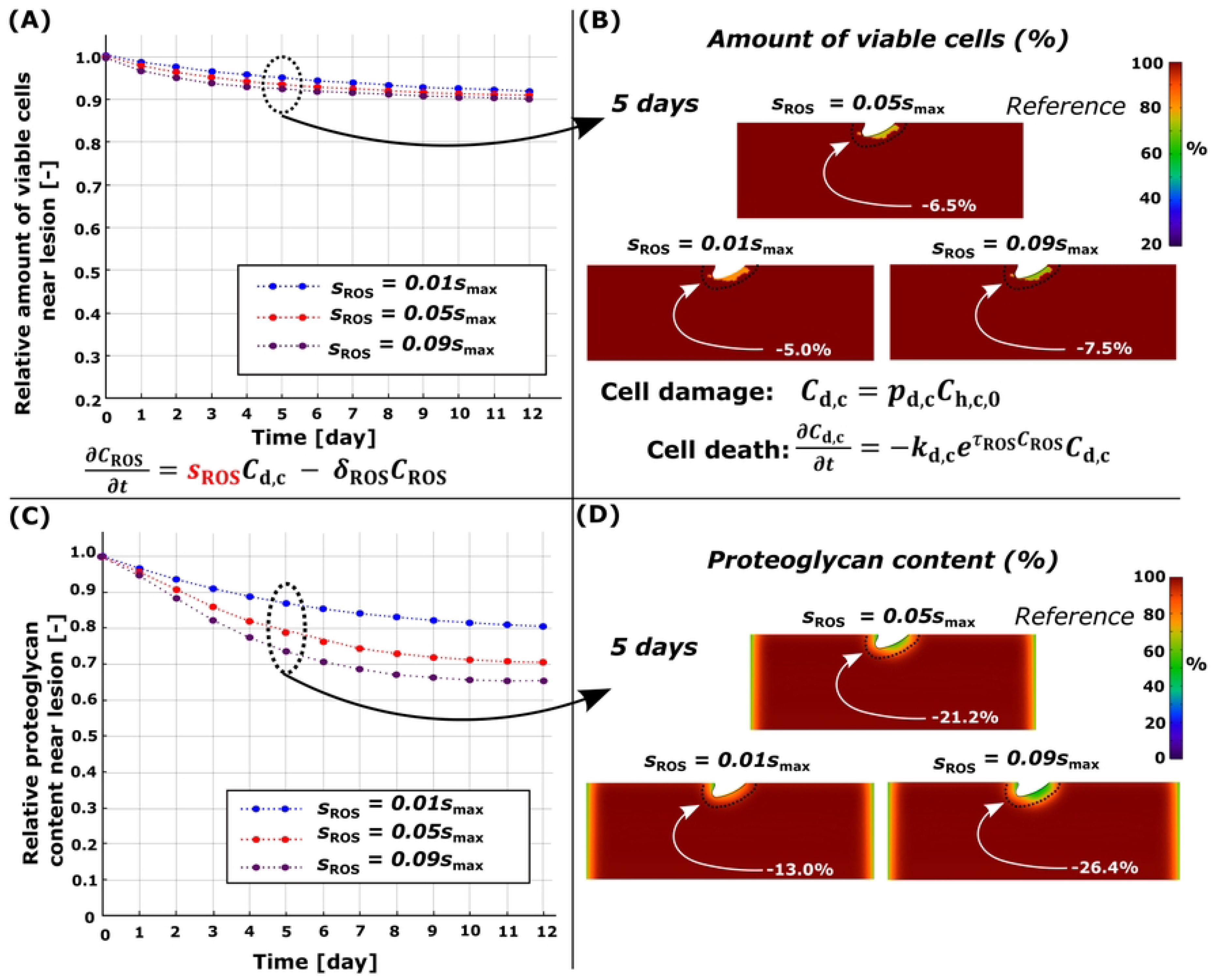
Sensitivity analysis of simulated reactive oxygen species (ROS) production rate *s*_ROS_. Comparison of temporal changes and spatial changes at day 5 in (A)-(B) in cell viability and (C)-(D) in proteoglycan content. (C) Higher simulated ROS production showed more intensive temporal proteoglycan loss and (A) cell death near the cartilage lesion compared to moderate and low production rates. Red line in (A) and (C) refers to the reference model (*s*_ROS_ = 0.40).

**Fig 6.**
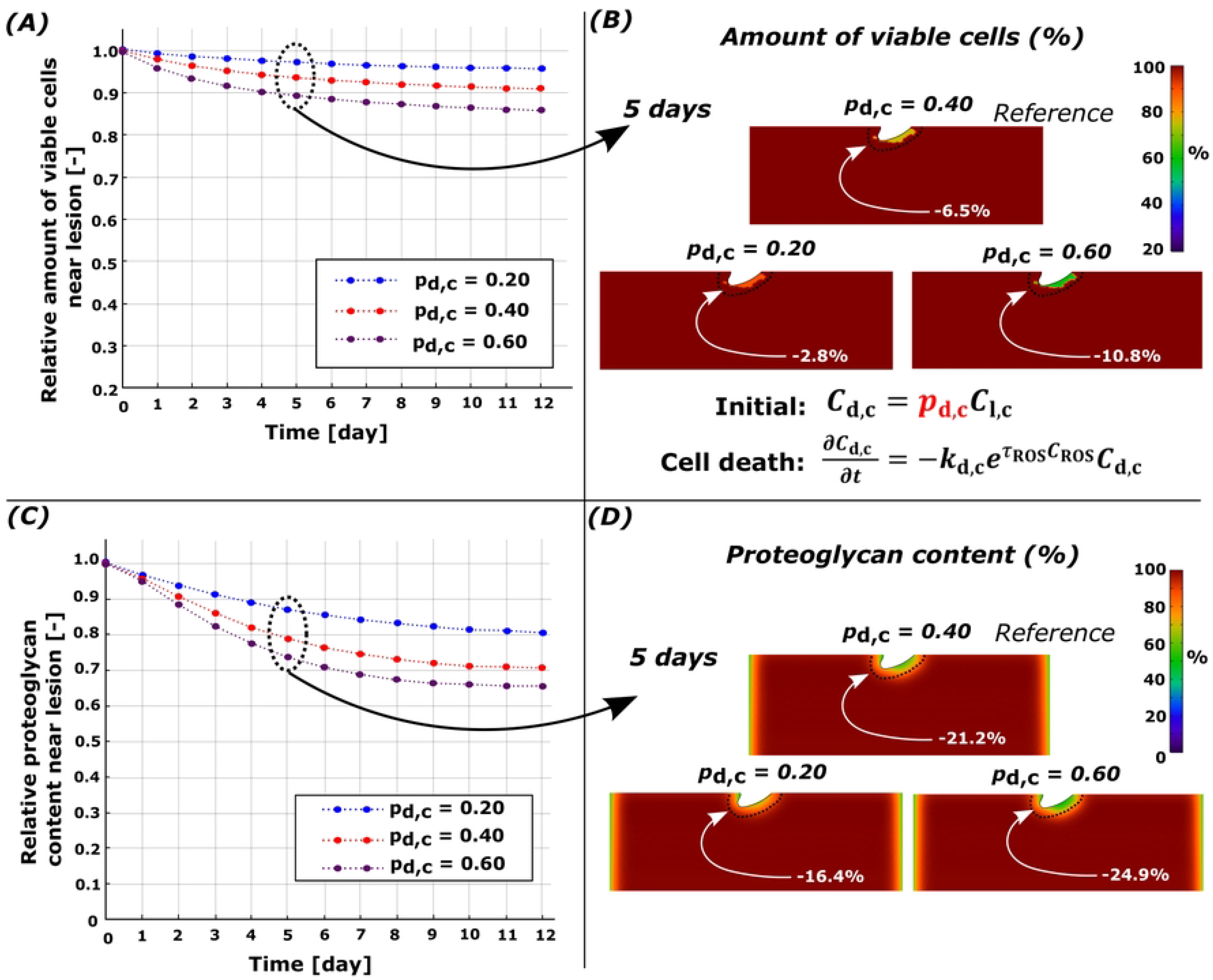
Sensitivity analysis for simulated damaged cell rate *p*_d,c_. Comparison of temporal changes and spatial changes at day 5 in (A)-(B) in cell viability and (C)-(D) in proteoglycan content. (A) Higher number of damaged cells led to more cell death and (C) more intensive proteoglycan degeneration near the cartilage lesion. Red line in (A) and (B) refers to the reference model (*p*_d,c_= 0.40).

### 3.3. Inflammation-induced apoptosis

Diffusion of IL-1 resulted in extensive cell apoptosis and subsequent PG loss near the free surfaces (Figs. 3C and 7). The model where proteoglycan degeneration via aggrecanases and loss of biosynthesis (induced by apoptosis) was considered showed PG loss of 50.4% near the cartilage lesion at day 5 (Fig. 3C). This rapid degradation masks the effect of IL-1 on PG loss through changes in PG biosynthesis. Thus, in Fig. 7, we present sensitivity analysis results with the effect of aggrecanases turned off in the model. At day 5, the reference model 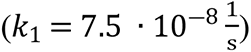 had PG loss of 11.2% (apoptosis of 33.5%) compared to day 0 (Fig. 7, red line). Corresponding models without apoptosis (*k*_1_ = 0) exhibited PG loss of 10.2% (Fig. 7, blue line; passive PG diffusion) and models with a higher apoptosis rate (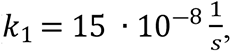 Fig. 7, purple line; apoptosis of 54.2%) showed PG loss of 11.9% in the cartilage.

**Fig 7.**
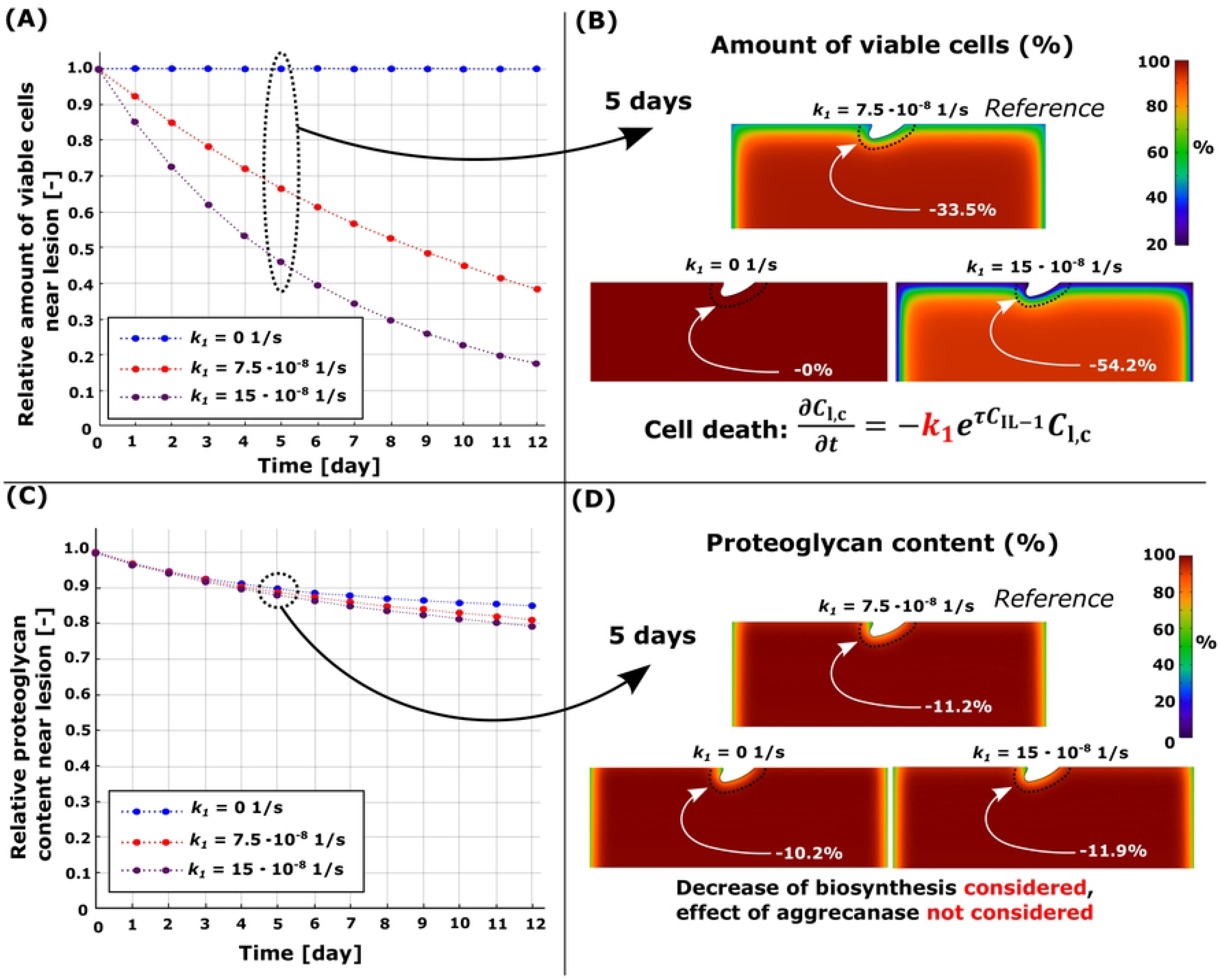
Sensitivity analysis for the simulated pro-inflammatory cytokine-induced apoptosis rate *k*_1_. Comparison of temporal changes and spatial changes at day 5 in (A)-(B) in cell viability and (C)-(D) in proteoglycan content. (A) Loss of viable cells and, thus, decrease of proteoglycan biosynthesis (aggrecanase induced proteoglycan degeneration was not considered), had (C) a negligible effect on the simulated proteoglycan content over 12 days. Red line in (A) and (B) refers to the reference model (*k*_1_ = 7.5 · 10^―8^1/s).

### 3.4. Synergistic effect of necrosis, ROS, and inflammation

Cartilage subjected simultaneously to the simulated effect of injury-related and inflammatory mechanisms revealed vast cell death and PG loss near the free surfaces and lesion (Figs. 3D and 8A-D). In the computational reference model (Fig 8A, red line), at day 5, near-lesion cell death was 46.8% (Fig 8C and D, total (bulk) cell death of 11.0% in the whole geometry) and PG loss was 64.2% (total PG loss of 18.9%) compared to day 0 (Fig 8E and F, red line). When the IL-1 concentration was decreased slowly in the combined model (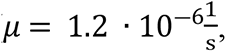, Fig. 8A, blue line), the simulated near-lesion cell death was 36.3% (Fig. 8C and D, blue line, total cell death of 8.1%) and PG loss was 62.0% (Fig. 8E and F, blue line, total PG loss of 16.7%). Rapid decrease (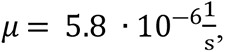, Fig. 8A, purple line) of IL-1 concentration in the culture medium led to near-lesion cell death of 25.6% (Fig. 8C and D, total cell death of 5.1% in the whole geometry) and PG loss of 50.8% (Fig. 8E and F, total PG loss of 10.9%). Interestingly, notably less PG loss was observed in 12-day simulations compared against the reference model (Fig. 8B).

**Fig 8.**
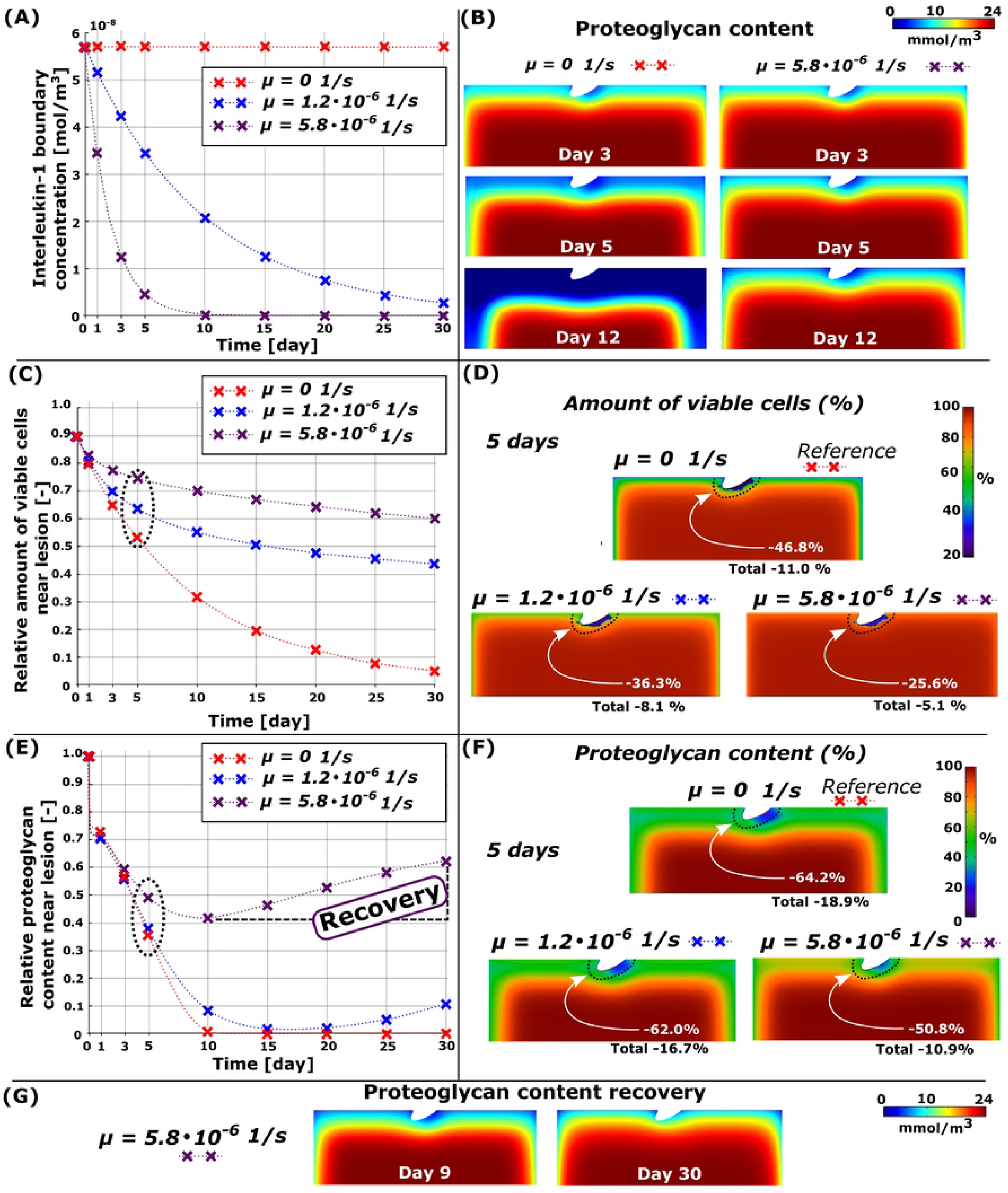
Simulated decrease of cytokine concentration in the combined model. (A) Simulated time-dependent exponential decrease of the interleukin-1 concentration in the culture medium and B) comparison of changes in proteoglycan (PG) content with constant (*μ* = 0 1/s) and fast-decreasing cytokine concentration (*μ* = 5.8 · 10^―6^ 1/s). C) Temporal changes in cell viability in 30-day simulation near the cartilage lesion (within 0.1 mm from the lesion) and D) spatial changes at day 5. D) Temporal changes PG content near the cartilage lesion and (F) spatial changes in the whole cartilage geometry (total) at day 5. Decreased exogenous cytokine concentration decreased cell death and PG loss substantially and (G) showed partial recovery of the PG content (here, simulation continued until day 30). Red line in the figure refers to the reference model (*k*_1_ = 1.2 · 10^―6^ 1/s).

### Partial recovery of the PG content in cartilage

When the simulation was continued until day 30, we observed that the greatest near-lesion PG loss of 98.5% and 58.2% occurred at day 17 and day 9 for the slow and fast decrease of IL-1 concentration. Moreover, we observed that at day 30, the PG content had recovered by 9.4% and 20.4% around the lesion (corresponding 4.0% and 3.9% bulk tissue recovery) for the slow and fast decrease of IL-1 concentration when compared to the PG content at days 17 and 9 (Fig 8G).

## 4. Discussion

Previous computational models of early PTOA have not explicitly modeled physical lesions, loading, or the underlying cell-regulated degradative mechanisms of cartilage. In this study, we bridged this knowledge gap and presented a novel mechanobiological model considering physical cartilage lesion, injury- and loading-related cell death, overproduction of ROS, and diffusion of pro-inflammatory cytokines. We compared the model results against previously measured optical density maps from injured calf cartilage samples and noticed matching predictions of the PG content: extensive and localized near the lesions, but more widely spread when IL-1 was added to the medium. Predicted cell death followed the same pattern of damage localization, observed also *in vitro*. The interesting computational findings are that 1) necrosis alone affects PG content rapidly (0–3 days) in the vicinity of the lesion but its effect almost completely fades away over 5 days, leading to partial recovery of PG content, 2) ROS overproduction and especially inflammation have longer-term (over 5 days) effects on PG content, and 3) rapid decrease of IL-1 concentration (leading to lower aggrecanase release and less suppression of PG biosynthesis) facilitates the recovery of PG content even in injured cartilage.

### 4.1. Necrosis

Injurious loading may cause rapid (within hours to days) necrotic and apoptotic cell death [12,13,17,54]. The injury can also sensitize live cells to turn catabolic more easily by later mechanical and inflammatory signals causing more extensive cell death if catabolic signals are not ceased [60, 61]. Locally elevated shear strains near lesions due to mechanical loading could be one such catabolic signal, assumed here to lead to localized necrosis and PG loss [16, 39]. The cell viability assay with propidium iodide and fluorescein acetate as used by Orozco et al. [16] and Eskelinen et al. [40] does not discern between necrosis and apoptosis, but other studies have shown that similar injurious loading may cause necrotic cell death [45]. Furthermore, the assumption that necrotic cells would release DAMPs inducing inflammatory response (such as IL-1 production, which later causes aggrecanase release [19]) is supported by several studies [20, 62].

On average, our model predicted necrotic cell death of 10.8% (40% local necrosis of the viable cells in areas exceeding 50% maximum shear strain threshold, Fig. 4A) within 0.10 mm from the lesion when collagen architecture was based on young bovine cartilage [16, 45]. For comparison, Philips et al. [45] reported a loss of cell viability around the superficial zone of mature bovine cartilage (0.15 ± 0.038 mm from the surface), especially in the vicinity of the surface fissures, 1 hour after impact-injury (unconfined compression with ∼25 MPa peak stress, 100%/s loading rate). Although we did not consider necrosis caused by the initial impact injury, our model predicted locally similar cell death near the lesion due to high strains resulting from dynamic loading of injured geometry, possibly causing rapid local degeneration of the PG content in the vicinity of the lesion.

In our simulated necrosis model, aggrecanases were released only at day 0 in response to cell necrosis near the lesion (Fig. 4A). We observed the substantial PG loss during the first 3 days near the lesion (Fig. 4C) and, as expected, simulating increased necrosis fraction led to higher PG loss, a scenario that is feasible with high impact loads [11,12,54]. Such an early local burst of enzymatic activity is supported by the finding that aggrecanase and other proteolytic enzyme expressions (e.g. MMP-3) increase within 1 day from experimental injury [8]. Moreover, studies about other arthropathies similar to osteoarthritis [21, 63] have suggested that the release of aggrecanases occurs in regions experiencing chondrocyte necrosis. Predicted PG degeneration within hours and the PG recovery within the following 3 days is explained by rapid outflux of aggrecanases from highly necrotic regions (change of aggrecanase concentration over time is relative to the aggrecanase concentration gradient) and relatively small effect of highly localized necrosis on total PG biosynthesis.

Interestingly, our results suggest that cartilage can recover its PG content partially and reach a steady-state after 12 days. This implies that after acute PG degradation and loss, decay of aggrecanase concentration and diffusion of synthesized PGs from deeper layers of the cartilage can promote PG recovery. However, in previous experiments [16, 40], PG degeneration was still observed near the lesion at day 12. This indicates that in addition to immediate necrosis, further mechanisms associated with cell damage (e.g. ROS overproduction) should be involved in the simulations to better catch the temporal changes in injured cartilage.

### 4.2. Cell damage, ROS, and apoptosis

Since maximum shear strains were excessive near the lesion, the damaged cells producing large amounts of ROS leading to apoptosis were located in the same areas as necrosis, with similarities to previous experiments with biological cartilage, where the amount of ROS was proportional to the deformation of the chondrocytes [49]. While the simulated necrosis indicated rapid early PG loss followed by partial PG content recovery, damaged cells contributing to the overproduction of ROS led to decreasing PG content over time. This suggests that necrosis might play an early short-term role in PG loss, but cell damage and its downstream catabolic effects may last longer despite the short lifetime of ROS [23, 54]. Thus, cell damage and large amounts of ROS could undermine the partial recovery seen with the necrosis model and continue cartilage degradation near the lesion even when tissue-level global loading is physiologically normal (15% strain in our model).

Low ROS production in cartilage did not result in major cell death (5.0%), nor did the moderate (6.5%) or severe (7.5%) ROS overproduction (Fig. 5B) near the lesion at day 5. Furthermore, low ROS production did not result in a substantial PG loss (13.0%, 2.8% higher than passive PG diffusion) whereas moderate and severe ROS overproduction resulted in higher PG loss, 21.2% and 26.4%, respectively. A similar interplay between damaged cells and increased ROS production leading to cell death and PG loss has been observed experimentally [23,49,53,54].

### 4.3. Inflammation

Simulated inflammation resulted in rapid and substantial cell death and PG loss near the free surfaces, in good agreement with experimental findings [29, 40]. At 1 ng/ml of IL-1, inflammation-driven degradation mechanisms dwarfed those driven by biomechanics. The inflammation-related PG loss was mostly driven by the aggrecanases; when the proteolytic effect of aggrecanases was turned off, the IL-1-induced apoptosis (resulting in decreased PG biosynthesis) had only a minor effect on the PG loss (Fig. 7C and D).

Analysis of inflammation-related PG loss has been extensively included in computational and experimental studies [29,34,36]. However, IL-1-induced cell death has rarely been included in computational models. In experimental work conducted by Lopez-Armada et al. [64], ∼50% bulk tissue cell death was observed after 7-day culture with 5 ng/ml of IL-1 [64], and Li et al. [29] reported ∼50% bulk cell death after 17 days culture with 1 ng/ml of IL-1. Our model exhibited 15.1% and 34.8% bulk cell death on days 7 and 17 with 1 ng/ml, respectively. Lower cell death in our simulated results could indicate that more inflammatory mechanisms such as pro-inflammatory cytokine and DAMP release from catabolic chondrocytes in cartilage [2, 18] are promoting cell death also in deeper layers of the cartilage, but this mechanism was omitted in our model.

### 4.4. Combined model

Simultaneously acting biomechanical and biochemical mechanisms resulted in marked cell death and PG loss especially near the lesion during the first 5 days (Fig. 3D and 8). Later, the IL-1-driven degradation dominated over the other mechanisms around the defect, in agreement with digital densitometry results [40]. Our model was able to capture the well-documented synergistic effect of biomechanics and inflammation on PTOA progression [61, 65].

Our reference model predicted locally extensive PG loss of 43.6% near the lesion at day 3 (Fig 8A and B, red line; total PG loss of 9.0% in the whole cartilage geometry at day 3) and spread of PG loss also to the intact areas at day 5 (Fig 8B and F; total PG loss of 18.9%). Eskelinen et al. [40] reported increased PG degeneration in the intact regions of injured-and-inflamed cartilage at day 7 compared to day 3. These experiments are in general consistent with our modeling results showing substantial near-lesion PG loss caused by synergistic effect of inflammation and high shear strains after 3 days and inflammation-induced PG loss also in the intact areas in the following time-points.

Interestingly, a simulated fast decrease of the IL-1 concentration in the culture medium resulted in partial recovery of the near-lesion (20.4% at day 30 compared to day 9) and bulk PG contents (3.9%). This finding highlights the major role of inflammation in the computational model; decreasing the cytokine concentration temporally leads to partial recovery of the tissue, while the biomechanical mechanisms contribute to tissue degradation around the lesions. The result of possible partial recovery suggests that inhibition of cytokine activity or rapid cytokine clearance from culture medium/joint space could suppress catabolic signals and bring cartilage closer to homeostasis.

### 4.5. Limitations

First, biomechanical loading and inflammation of cartilage include many complex cell-level mechanisms. Although our approach represents a step toward elucidating the degradation mechanisms after injury, our model has limitations that may partly explain the disagreement between the model and experiments. Additional degenerative mechanisms to consider are the IL-1-induced ROS production [52], ROS-induced necrotic cell death [15, 66], the introduction of MMP-3-driven matrix degradation after injury [11, 67], fluid flow-dependent PG loss through lesion edges [16], injury-related PG loss due to microdamage and structural changes instead of enzymatic degradation [60], and beneficial effects of moderate cyclic loading [68]. Furthermore, we used the simulated IL-1 concentration of 1 ng/ml, the same as used in previous experimental in vitro studies [29,34,36]. After acute inflammation, physiological IL-1 concentration in the inflamed knee joint is typically much lower than 1 ng/ml [32, 69], but the model was calibrated previously based on in vitro experiments, and the use of physiological concentrations would just result in slower progress of the degeneration. Furthermore, we did not account for the degeneration of the collagen network that would affect the biomechanical properties and cell responses in the cartilage [70]. This was justified as structural and constitutional changes in the collagen network have been observed to occur later than those of in the PG content [29, 71]. Also, we did not explicitly consider the pericellular matrix or changes in its properties during the degeneration. There is evidence that alterations in the pericellular matrix properties and cell-matrix interactions may have substantial role in the OA initiation and progression of tissue degeneration [72–74], thus, the function of the pericellular matrix should be accounted for in future studies.

Second, the biomechanical loading used in the computational model is a simplification of the experiments. For instance, we did not simulate the initial impact-loading leading to cartilage defects in the superficial zone or the full cyclic loading protocol used in previous experiments after the injury [16, 40]. We note that the compositional changes after impact-loading and during the continuous cyclic loading can influence the shear strain distributions, leading to more severe cartilage degeneration than currently predicted by the model [75].

Third, although some inflammation and material-related parameters have been well-calibrated [16, 36], model validity testing is hampered by the small amount of biomechanical and biochemical experimental data, which are only available at a few time points. Therefore, the model has several biochemical parameters that need to be better calibrated. For example, the parameters for ROS production, necrotic/damaged cell fraction, and possible necrosis/apoptosis-related release of aggrecanases [4,21,63] and matrix degradation require further experimental support. However, the presented predictions are already generally in line with the current literature and despite the lack of extensive calibration, the current modeling framework can offer insights into the mechanisms driving cell death and proteoglycan loss in PTOA-like conditions.

### 4.6. Future directions

In the future, multiscale mechanobiological models may be a feasible pathway to produce patient-specific predictions of early cartilage degeneration, open new avenues for high-level translational research and be a tool to assess different intervention strategies to mitigate PTOA progression. However, extensive experimental research is still needed to elucidate the injury-related mechanotransduction pathways, cell death, and ROS kinetics, which could provide time-dependent quantitative data to calibrate and enhance current models. Specifically, interesting future aims are to include the function of the pericellular matrix and sensitization of near-lesion cells to further damage if biomechanical or inflammatory challenge is not removed [61], and to model the effects of ROS-suppressing disease-modifying drugs [7, 54]. Ultimately, the calibrated mechanobiological cell-tissue-level models could be augmented to the joint level, which could be used to produce cost-efficient optimized intervention strategies to mitigate early cartilage degeneration.

## 5. Conclusions

Cell death and enzymatic cartilage degeneration in response to injurious loading are important factors to consider in computational models for predicting PTOA progression. We incorporated biological cell–tissue-level responses including necrotic and apoptotic cell death, ROS overproduction, and inflammation of injured cartilage into a finite element model of early-stage PTOA. Our novel mechanobiological model was able to predict localized cell death and PG loss similar to previous biological experiments; biomechanically induced necrosis and apoptosis and the following enzymatic degeneration of PGs were observed near the cartilage lesion, while diffusing pro-inflammatory cytokines resulted in more widely spread damage. Based on the computational model predictions, rapid inhibition or clearance of pro-inflammatory cytokines would result in partial recovery of the PG content and could be a potential way to decelerate PTOA progression even in injured tissue. In the future, the current computational framework could enhance previous models by introducing new mechanisms, thus providing a better understanding of PTOA progression. Furthermore, in the future, thoroughly calibrated multi-level mechanobiological models could be a valuable tool in assessing patient-specific pharmacological treatments time-dependently and help in the planning of new more efficient intervention strategies.

## Acknowledgements

We acknowledge the support of University of Eastern Finland, Massachusetts Institute of Technology and University of Iowa to conduct this research.

## Supporting information captions

**S1 Biomechanical material model.** The supplementary material providing more detailed information about the biomechanical material model.

**S1 Table. Variables describing cartilage composition**. Normalized depth *z* is defined as *z* = 0 on the injured surface and *z* = 1 on the bottom surface.

**S2 Boundary conditions and finite element mesh used in biomechanical simulations.** The supplementary material providing more detailed information about the simulations and boundary conditions of the biomechanical model.

**S2 Fig. Finite element mesh used in the biomechanical simulations.** Finite element mesh for the injured cartilage geometry including 918 linear axisymmetric elements with pore pressure.

**S3 Higher axial strain amplitude to investigate the initial impact injury.** The supplementary material providing additional analysis on the injurious loading used in the experiments.

**S3 Fig. Additional analysis of the higher axial strain to study injurious loading.** Finite element mesh for the intact geometry and the maximum shear strain distributions after unconfined compression with 40% axial strain amplitude.

**S4 Reaction–diffusion model: simulated changes in proteoglycan content.** The supplementary material providing more detailed information about the biochemical reaction–diffusion model.

**S5 Mesh sensitivity.** The supplementary material providing more detailed information about the mesh sensitivity analysis.

**S5 Fig. Mesh sensitivity analysis.** Mesh sensitivity analysis for the mechanobiological simulations conducted with the combined model.

**S6 Data interpolation to Comsol.** The supplementary material providing more detailed information about the data interpolation from the biomechanical simulations.

